# Structural connectivity of auditory-linguistic brain networks predicts success in speech categorization and listening in noise

**DOI:** 10.64898/2026.07.06.736789

**Authors:** Rose Rizzi, Jack R. Stirn, Zara Eisenhut, Gavin M. Bidelman

## Abstract

Successful speech perception requires listeners to bin continuous acoustic information into discrete phonetic categories. However, some people maintain within-category acoustic information (gradient) while others discard category-irrelevant information (discrete) during perception. Listeners also vary in how consistently they label speech sounds and more gradient/consistent labeling has been linked with better speech-in-noise (SIN) perception. Here, we test how neuroanatomical properties of the brain’s major speech-language and auditory pathways relate to individual differences in speech categorization and SIN processing. We measured phonetic categorization and SIN comprehension via phoneme labeling and QuickSIN tasks. Diffusion-weighted imaging (DWI) with probabilistic tractography estimated axonal density within the bilateral arcuate fasciculi and brainstem-cortical auditory projections. Anatomical morphology (surface area, gray matter volume, thickness) was also quantified in the adjacent frontotemporal cortical areas and midbrain. Behaviorally, we found more consistent categorizers had better performance on the QuickSIN. DWI showed that more gradient listeners had greater white matter density in the left arcuate fasciculus and brainstem-cortical auditory pathways, while better SIN performance was predicted by denser white matter in the brainstem-cortical auditory pathways. Morphometric results revealed more consistent listening was associated with greater cortical thickness in right superior temporal gyrus and more gradient listening was associated with greater surface area in right pars opercularis. We infer that individual differences in phonetic categorization relate to SIN comprehension and are at least partially explained by neuroanatomical properties of the auditory-linguistic brain.

## 1. Introduction

Listeners differ in how they categorize speech sounds. They vary in how gradiently vs. discretely they label speech sounds (Kapnoula et al., 2021; Kapnoula et al., 2017; Kong & Edwards, 2016; Myers et al., 2024; Rizzi & Bidelman, 2024) and in how they weigh within-vs. between-category information to inform their perceptual identification (Kapnoula et al., 2017; McMurray, 2022; Rizzi & Bidelman, 2024). A more gradient listener uses fine-grained acoustic information, while a more discrete listener more heavily weights abstract categorical representations. This weighting results in more gradient listeners having a more linear mapping of acoustics to perception and more discrete listeners having a perceptual space that is strongly warped onto category labels. The degree of perceptual gradience in a listener’s responses can be quantified as the slope of their identification curves derived from labeling speech sounds along an acoustic-phonetic continuum. As in most categorical perception experiments, shallower slopes represent more gradient listening and steeper slopes represent more discrete listening (e.g., Bidelman et al., 2026a; Kapnoula et al., 2021; Kapnoula et al., 2017; Rizzi & Bidelman, 2024).

Similarly, listeners can vary in how consistently they categorize speech sounds (Kim et al., 2025a; Kim et al., 2025b; Kim et al., 2024; Rizzi & Bidelman, 2025). For example, some listeners choose the same label for a speech sound across trials, while others might use different labels across repeated presentations of the same token. Typical categorical perception task paradigms using a two-alternative forced choice (2AFC) task confound the perceptual properties of gradience and consistency: a shallow identification slope could derive from a consistent gradient listener or an inconsistent discrete listener. Moreover, consistency cannot be directly measured with a 2AFC because the nature of the binary response imposed by a 2AFC conflates gradience with consistency (Apfelbaum et al., 2022; Kapnoula et al., 2017; McMurray, 2022). A listener with a shallower identification curve slope from a 2AFC could either be a more gradient or a noisier (less consistent) responder, resulting in averaged responses that are closer to chance. However, consistency and gradience can both be assessed during phoneme labeling with a visual analog scale (VAS) paradigm which avoid binary labels and allows more graded responding to an acoustic-phonetic continuum (Apfelbaum et al., 2022). Thus, individual differences in speech categorization can be more comprehensively assessed when listeners label speech tokens from an acoustic-phonetic continuum using a VAS.

It has been hypothesized that gradience and consistency in auditory categorization might also relate to other important listening skills, mainly speech-in-noise (SIN) processing. If more gradient listeners have more flexible perception and better recovery from ambiguity (Kapnoula et al., 2021; Kapnoula et al., 2017; Kutlu et al., 2024), they may be better able to adapt to challenges posed by noise-degraded listening. Findings investigating this hypothesis have been somewhat mixed, with some studies finding gradient listening benefits SIN perception (Myers et al., 2024; Rizzi & Bidelman, 2024) while others fail to observe a relationship between gradience and SIN skills (Kapnoula et al., 2021; Kapnoula et al., 2017; Rizzi & Bidelman, 2025). These mixed findings may be due to high inter-subject variability in gradience especially when assessed across different phonetic contrasts and at the word and sentence level (Kim et al., 2025b; Myers et al., 2024). In contrast, consistency seems to be a more trait-like index of perceptual categorization that remains stable within listener even when assessed across varied phonetic contrasts (Kim et al., 2025b). Consistency may also be a more sensitive measure of between-subject differences in categorization as it can measure subtle trial-by-trial variance in a listeners’ responses that are lost when averaging data across many trials (Kim et al., 2025b). Thus, if a relationship does exist between categorization skills and SIN perception, consistency may be a stronger, more stable predictor of SIN understanding than gradience (Myers et al., 2024; Rizzi & Bidelman, 2025). More consistent listeners might have a more robust perceptual readout that is less degraded by noise than in an inconsistent listener. Using EEG, we have recently shown that perceptual consistency is related to increased neural consistency in how the brain encodes speech sounds on a trial-by-trial basis (Rizzi et al., 2026). This provides a functional mechanism to account for differences in behavioral categorization. More consistent categorization has also been linked to improved SIN performance (Myers et al., 2024; Rizzi & Bidelman, 2025; Rizzi et al., 2026). While our other work has examined *functional/electrophysiological* correlates of perceptual gradience and consistency (Rizzi & Bidelman, 2024; Rizzi et al., 2026), it is unclear how underlying anatomical brain structures or pathways might also relate to these perceptual properties of speech categorization.

In this vein, neuroimaging studies have revealed that phonetic categories are clearly represented in inferior frontal gyrus (IFG) (Binder et al., 2004; Blumstein et al., 2005; Lee et al., 2012; Luthra et al., 2019; Myers, 2007; Myers et al., 2009; Rogers & Davis, 2017; Toscano et al., 2018). However, some studies have observed categorical organization in neural responses even earlier in the neural hierarchy in superior temporal gyrus (STG) (Bidelman & Walker, 2019; Chang et al., 2010; Desai et al., 2008; Liebenthal et al., 2005; Mesgarani et al., 2014) and auditory brainstem (Carter & Bidelman, 2023; Rizzi & Bidelman, 2023). Neural responses to speech become more abstract along the auditory pathway, with some neurons selectively responding to speech over nonspeech in STG (Benson et al., 2001; Binder et al., 2000; Humphries et al., 2014; Zatorre et al., 1992) but not primary auditory cortex (Hamilton et al., 2021). And within IFG, the pars opercularis and triangularis are particularly involved in categorical computations (Lee et al., 2012; Mahmud et al., 2021; Papoutsi et al., 2009). Such findings are consistent with the idea that phonetic categories are likely coded at higher levels of the auditory system and downstream from initial sound arrival in A1. While categorical representations are observable in human brainstem FFRs, they have only been observed under attentive states, suggesting category coding is not intrinsic to brainstem, per se, but rather inherited from top-down corticofugal tuning of subcortical activity from higher cortical structures (Bidelman et al., 2013; Carter & Bidelman, 2023; Rizzi & Bidelman, 2023). Together, it is clear that a network of auditory and linguistic brain regions including brainstem, STG, and IFG are relevant to acoustic-phonetic mapping in speech perception.

To date, few studies have investigated relationships between the morphology of auditory-linguistic brain *structures*, the white matter tracts connecting them, and variance in categorization gradience and consistency. Notably, Fuhrmeister and Myers (2021) found reduced surface area in right middle frontal gyrus predicted more gradient fricative labeling and increased gyrification of bilateral transverse temporal gyri predicted less consistent speech categorization, supporting the notion that frontal regions are sensitive to categorical information. While Fuhrmeister and Myers (2021) used a VAS to assess phoneme labeling, the task used discretized steps which provided 7 anchor points (with a 7-token continuum) for listeners to place their response. This design makes it difficult to differentiate between response profiles of listeners attempting to bin tokens into smaller, yet discrete categories, and those utilizing a more continuous listening strategy. Likewise, a listener may appear more consistent under this task if they have anchors to guide their responses. Thus, the structure-behavior relations observed in Fuhrmeister and Myers (2021) could be confounded. In the present study, we use a fully continuous response paradigm to better elucidate the relationships between structural properties of the brain’s auditory-linguistic pathways and perception gradience and consistency.

While volumetric findings in Fuhrmeister and Myers (2021) were restricted to surface area and local gyrification, other metrics of anatomical morphology can also be extracted from MRI brain images including cortical thickness and gray matter volume. There are subtle differences in these measures, with surface area reflecting the number of cell columns in the cortical region and thickness reflecting the number and density of cells within those columns (Rakic, 1988). Gray matter volume is the combination of thickness and surface area, representing both the number of cellular columns and the number of cells within those columns. There are distinct genetic bases (Panizzon et al., 2009; Winkler et al., 2010) and developmental trajectories (Wierenga et al., 2014) for cortical thickness and surface area, suggesting they could differentially relate to behavior. Furthermore, cortical thickness and surface area are phenotypically independent both globally and regionally (Panizzon et al., 2009; Winkler et al., 2010), emphasizing they may have distinct contributions to perception. Thus, measuring all three metrics can aid in a deeper understanding of structure-behavior relationships.

While studies show that fronto-temporal brain regions are critically involved in categorization (e.g., Alho et al., 2016; Bidelman & Walker, 2019; Blumstein et al., 2005; Husain et al., 2006b; Lee et al., 2012; Myers et al., 2009), none to our knowledge have explicitly investigated how connectivity between these structures relate to phonetic categorization skills. White matter (WM) tract density can be readily measured using diffusion weighted imaging (DWI), a form of magnetic resonance imaging (MRI) that measures the amount of water diffusion in the brain in each cardinal direction. In general, water molecules move more easily within and along than through axons. DWI scans can then be used with probabilistic tractography algorithms to construct a map of the major fiber tracks in the brain. From fiber tracking, different metrics of WM density and integrity, such as quantitative anisotropy (QA), can be estimated to understand the microstructural properties of the neuroanatomy. QA provides an estimate of WM density that is more robust to crossing fiber tracks than some other diffusion metrics (Shen et al., 2015; Yeh et al., 2013).

Given the role of STG and IFG in categorization, it is plausible that WM density of arcuate fasciculus (AF) could relate to phonetic categorization. The classical (direct) AF pathway connects Wernicke’s (posterior STG) and Broca’s (IFG) areas with an additional indirect pathway running through the parietal lobe (Catani et al., 2005). Specifically, the indirect pathway can be divided into an anterior segment connecting Broca’s area to the inferior parietal lobe and a posterior segment connecting the inferior parietal lobe to Wernicke’s area (Catani et al., 2005). The first reported function of the AF was for repeating verbal information, an ability that is impaired in conduction aphasia (Wernicke, 1874). However, more modern views acknowledge broader functions of the AF. In the dual stream model of speech, the AF provides direct anatomical connection for the dorsal stream which supports the mapping of acoustic signals to frontal articulatory networks (Hickok & Poeppel, 2007). This dorsal stream is largely left-lateralized and underlies sub-lexical speech sound segmentation and short-term phonological memory and guides articulation, while the ventral stream plays a larger role in comprehension (Hickok & Poeppel, 2004, 2007). Though not originally proposed as a direct function of the dorsal stream, functional neuroimaging studies have revealed patterns of connectivity suggesting phonetic categorization relies on the dorsal stream (Chevillet et al., 2013; Rauschecker, 2012; Zaehle et al., 2008). Thus, commonly observed functional activations of IFG and STG observed in categorization studies likely involve the AF. Supporting this notion, studies have demonstrated microstructure of the AF is related to sensitivity (Perron et al., 2021; Tremblay et al., 2019) and accuracy (Li et al., 2021) of CV discrimination in noise. Microstructure of the AF also relates to phonological processing and dyslexia in left hemisphere (Aeby et al., 2013; Langer et al., 2015; Lebel & Beaulieu, 2009; Perdue et al., 2025; Vandermosten et al., 2012; Yeatman et al., 2011) and to pitch perception in right hemisphere (Chen et al., 2018; Loui et al., 2009; Loui et al., 2011). Thus, it is plausible the AF similarly supports the conversion of acoustic to phonetic information during speech categorization.

WM microstructure along the canonical auditory pathway connecting lower processing centers (brainstem) to primary auditory cortex could also relate to speech processing and categorization. However, few studies have examined white matter tractography of the lemniscal auditory pathways. Among the handful of reports, declines in WM of the brainstem-cortical projections have been linked to tinnitus and hearing loss (Koops et al., 2021; Svobodová et al., 2024). Though it is plausible that denser WM bundles in the central auditory pathways might provide a higher fidelity readout to support speech perception in noise or retain subphonemic acoustic detail, no studies have assessed whether brainstem-cortical WM microstructure relates to auditory perception. Moreover, how categorization gradience/consistency and SIN perception relate to language (AF) pathways also remains unknown. If categorization relates more strongly to auditory system anatomy, this would suggest that acoustic-to-phonetic mapping is a somewhat automatic process and depends on integrity of low-level hearing pathways. On the contrary, if categorization more strongly relates to language pathway anatomy (AF), this would argue that acoustic-to-phonetic processing requires later control mechanisms downstream from the canonical auditory system. Lastly, if categorization relates to microstructure of both auditory and language WM tracts, it may rely on an interplay between the strength of early auditory encoding and later involvement of auditory-motor integration as implied by the dual stream theory of speech-language processing (Hickok & Poeppel, 2004).

Here, we extended the functional results of our prior work (Rizzi & Bidelman, 2024; Rizzi et al., 2026) by investigating whether there is also a *structural basis* to categorization skills (gradience and consistency) and SIN processing. We used MRI volumetrics and DWI neuroimaging to quantify the anatomical morphology and microstructure connectivity within and between major auditory and language regions in cortex (STG and IFG subdivisions) and the subcortical auditory pathways. We hypothesized that more discrete listeners would have larger regional volumetrics in IFG, while more consistent listeners would have larger volumetrics in STG. We further hypothesized that density of both AF and auditory brainstem-cortical fiber tracts would relate to variance in categorization skills, emphasizing the role of both the auditory system and later integration via the dorsal stream for phonetic categorization. Specifically, we expected that more gradient listeners who have greater sensitivity to fine acoustic-phonetic details would have denser brainstem-cortical projections and AF, as implied by previous studies (Perron et al., 2021; Tremblay et al., 2019). Finally, we hypothesized that denser WM in both auditory and language pathways would also relate to better SIN processing, suggesting an anatomical mechanism for the relationship between categorization and SIN ability.

## 2. Materials and Methods

The sample included a subset of listeners from our companion EEG experiments who had no contraindications for MRI scanning (EEG data are reported in Rizzi et al., 2026). This included *N* = 31 young adults (22.65 ± 4.75 years; 9 male, 22 female) with 16.32 ± 2.77 years of education and 8.10 ± 5.96 years of formal music training. Participants were mostly right-handed (75% ± 41% Edinburgh Handedness Inventory; Oldfield, 1971). All participants had normal hearing and had English as their first language. Participants provided written informed consent in accordance with a protocol approved by the Institutional Review Board at Indiana University and were paid $15 an hour for their time.

### 2.1 Behavioral measures

Behavioral data were collected as described in Rizzi et al. (2026). Listeners labeled 100 ms vowels along an acoustic-phonetic continuum from /u/-/a/ (for acoustic details, see Bidelman et al., 2013; Rizzi & Bidelman, 2024, 2025) using a visual analog scale (VAS). Phoneme labeling occurred during EEG recording in the first lab visit. We fit individual subject identification curves with a sigmoid P = 1/[1 + e^−β1(x−β0)^] and estimated parameters using the *psignifit* function (Schütt et al., 2016) in MATLAB (v2024a). To quantify gradience, we estimated the psychometric slope (β*1*) from VAS labeling during EEG recording. Consistency was quantified as 1 – σ of VAS responses (distance clicked along the scale). We assessed SIN ability for each listener using the average SNR Loss from 2-4 lists of the QuickSIN (Killion et al., 2004).

### 2.2 MRI

We collected 3D whole-brain T1-weighted anatomical volumes from each participant (MPRAGE; TE/TR=2.7 ms/2400 ms, 160 axial slices, voxel size=1×1×1 mm3, slice thickness =0.8 mm; FOV=256mm, FA=80) using the 3T Siemens Magnetom Prisma scanner housed in the IU Imaging Research Facility. MRI data were converted to ezBIDS format (https://brainlife.io/ezbids/; Levitas et al., 2024).

### 2.3 Segmentation

We used the FreeSurfer (v7.3.1; http://surfer.nmr.mgh.harvard.edu/) comprehensive *recon-all* pipeline (Fischl, 2012) to perform cortical reconstruction and volumetric segmentation on the T1-weighted structural MRI scans using. This automated pipeline includes motion correction (Reuter et al., 2010), removal of non-brain tissue and skull stripping via hybrid watershed/surface deformation (Fischl et al., 2004), automated Talairach transformation, segmentation of white and gray matter volumetric structures (Desikan et al., 2006; Fischl et al., 2004), and cortical surface reconstruction (Dale et al., 1999). MRI processing was conducted on the Indiana University high-throughput Quartz supercomputing cluster (92 compute nodes, each equipped with two 64-core AMD EPYC 7742 2.25 GHz CPUs and 512 GB of RAM).

From the full-brain FreeSurfer output (i.e., aparc + aseg stats table from *recon-all*), we measured cortical thickness (mm), surface area (mm^2^), and gray matter volume (mm^3^) (Dale et al., 1999; Fischl & Dale, 2000; Fischl et al., 1999; Fischl et al., 2004) from each region defined in the Desikan-Killiany atlas parcellation (Desikan et al., 2006). FreeSurfer morphometrics have good test-retest reliability across scanners and various field strengths (Han et al., 2006; Reuter et al., 2012). ROIs for analysis included the six regions needed to test our hypothesis: bilateral superior temporal gyrus (STG) and bilateral pars opercularis (PO) and pars triangularis (PT) of the IFG. We did not analyze brainstem volume since other metrics (thickness, surface area) cannot be computed from the subcortical atlas.

### 2.4 Diffusion weighted imaging (DWI)

We used a multishell DWI scheme with two runs of DWI acquisitions with opposite phase-encoding directions (TR/TE=3516/88 ms, SMS acceleration factor=4, b-values=1000 and 2500 s/mm^2^, sampling directions=76 and 74, plus 5 volumes with b=0, isotropic in plane resolution=1.5 x 1.5 x 1.5 mm^3^ with whole-brain coverage, total scan time=10 mins). The two DWI scans were combined and corrected for susceptibility and Eddy current distortions using the FSL software suite (Jenkinson et al., 2012). Deterministic fiber tracking was then performed using DSI Studio software (version “Chen”) (https://dsi-studio.labsolver.org/) (Yeh et al., 2016; Yeh et al., 2010). The diffusion data were reconstructed in the MNI space using q-space diffeomorphic reconstruction (Yeh & Tseng, 2011) to obtain the spin distribution function (Yeh et al., 2010). A diffusion sampling length ratio of 1.25 was used. The output resolution in diffeomorphic reconstruction was 1.5 mm isotropic. The restricted diffusion was quantified using restricted diffusion imaging (Yeh et al., 2017). The tensor metrics were calculated using DWI with b-value lower than 1750 s/mm². There was one scan with poor quality that was excluded from further analysis, resulting in 30 total scans.

### 2.5 Fiber tracking

We computed fiber tracks restricted to the bilateral auditory brainstem-cortical pathways based on the HCP tractography atlas as defined in DSI studio (Yeh et al., 2018) and ICBM 152 adult template brain landmarks (Mazziotta et al., 1995). Auditory tracks were generated by seeding ROIs in the “brainstem” and “auditory sensory” regions of the CerebrA and Campbell atlases, respectively. This constraint produced similar streamlines to the in vivo human auditory brainstem white matter atlas described by Sitek et al. (2019). Autotrack was used to automatically identify auditory fibers with a distance tolerance of 24 mm in the ICBM152 space. The anisotropy threshold was randomly selected between 0.5 and 0.7 Otsu threshold. The angular threshold was randomly selected from 45 to 90°. The step size was set to voxel spacing. A total of up to 5000 streamlines were generated for each pathway. Auditory tracks with length shorter than 5 or longer than 100 mm were discarded. We also computed tracks restricted to bilateral arcuate fasciculus. AF tracks with lengths shorter than 30 mm or longer than 300 mm were discarded. Autotolerance for tracking was set at 24. We then extracted the mean quantitative anisotropy (QA) of each tract bundle per participant, reflecting the overall density of each anatomical pathway. Left and right hemispheres were analyzed separately.

### 2.6 Statistical Analysis

Unless otherwise specified, we used linear mixed effects models (lme4 package version 1.1-32 in R version 4.2.1; Bates et al., 2015) to assess differences in neural (MRI volumetrics and WM tract QA) and behavioral (categorization consistency, gradience, and QuickSIN score) dependent variables. Full model details are reported below. Pairwise contrasts were Tukey-adjusted to account for multiple comparisons and family-wise error corrections were employed where appropriate. Degrees of freedom for mixed models were estimated using Satterthwaite’s method. Handedness (*p* > 0.85) and years of music training (*p* > 0.05) did not relate to behavioral measures in our sample and were therefore not included in our statistical models.

## 3. Results

### 3.1 Behavioral data

Categorization consistency and SIN perception were correlated [*r*(28) = -0.39, *p* = 0.038]. More consistent categorizers had better SIN scores relative to inconsistent categorizers (**Fig. 1**).

**Figure 1.**
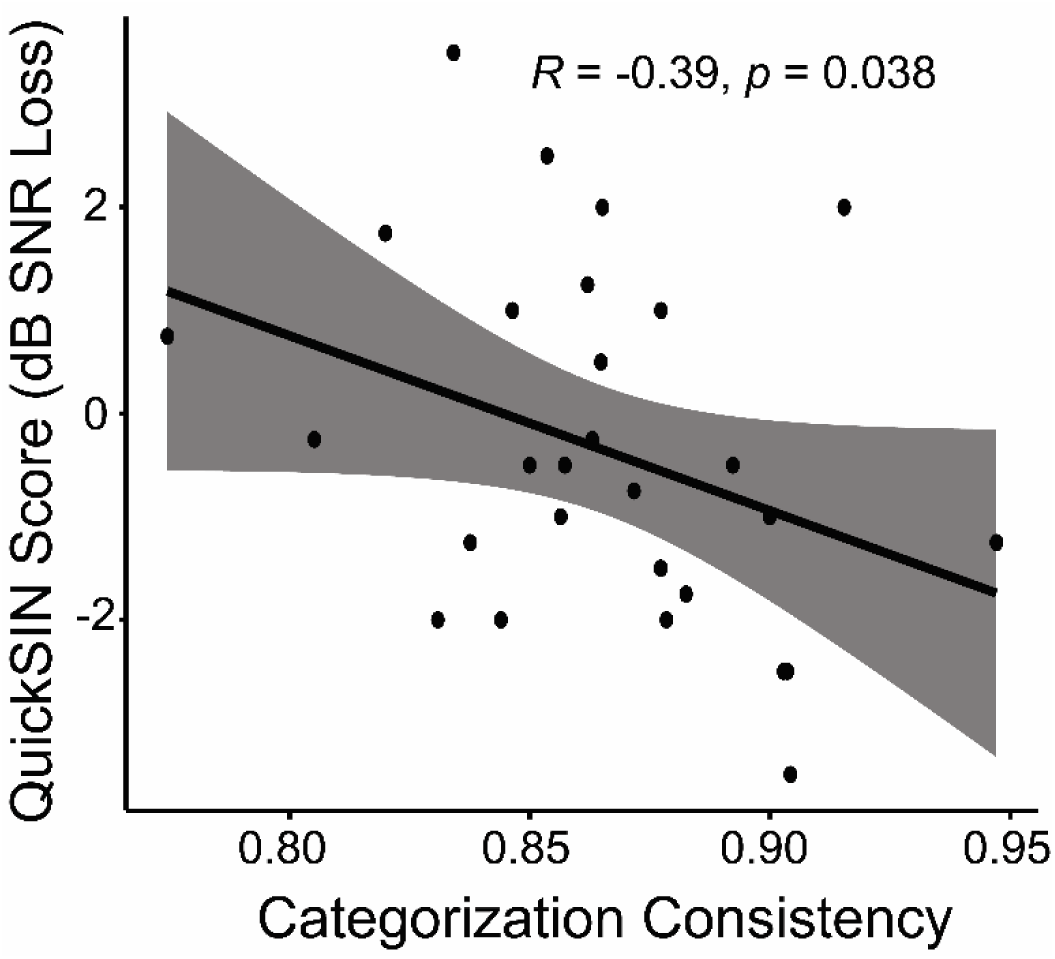
More consistent categorizers in vowel labeling perform better on the QuickSIN. Shading = 95% CI.

### 3.2 Structural data

Surface area, gray matter volume, and cortical thickness of auditory-linguistic ROIs are shown in **Fig. 2**. We built linear mixed effects models with each MRI measure as the outcome with a fixed effect of ROI and random slopes for subjects. Surface area [*F*(5, 140) = 906.49, *p* < 0.0001, *η^2^_p_* = 0.97], gray matter volume [*F*(5, 140) = 819.94, *p* < 0.0001, *η^2^_p_* = 0.97], and cortical thickness [*F*(5, 140) = 102.21, *p* < 0.0001, *η^2^_p_* = 0.78] all expectedly varied across ROIs. Tukey-adjusted pairwise comparisons revealed surface area (SA) and gray matter volume (GMV) were larger in LH STG (*p_SA_* = 0.0001; *p_GMV_ =* 0.004) and pars opercularis (PO) (*p_SA_*= 0.0001; *p_GMV_ =* 0.0008) relative to their RH counterparts (see **Table 1**). Pars triangularis (PT) SA and GMV were similar between hemispheres (both *p* > 0.31). Gray matter volume and surface area were larger in left hemisphere PO than PT (*p_SA_ =* 0.0001; *p_GMV_*< 0.0001) and were invariant in the right hemisphere (both *p* > 0.319). Cortical thickness (CT) varied across ROIs (*p*s < 0.0001) but was similar between hemispheres (*p* > 0.83).

**Figure 2.**
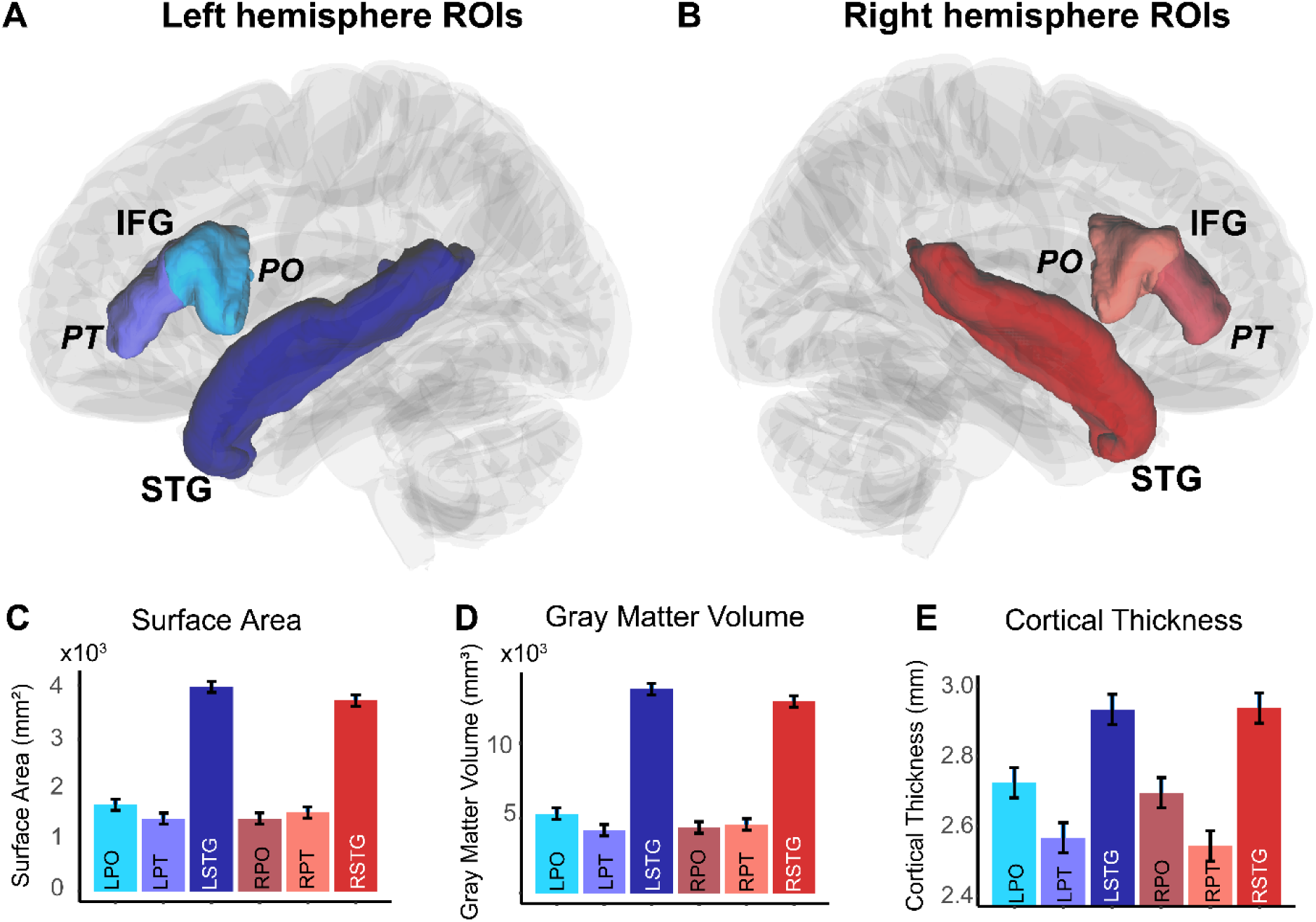
Structural morphology of auditory-linguistic cortical brain regions. (A-B) Bilateral ROIs from the Desikan-Killiany atlas. Superior temporal gyrus (STG): dark blue and red. Inferior frontal gyrus (IFG): pars triangularis (PT - anterior), pars opercularis (PO - posterior) in lighter blue and red. (C-E) MRI structural estimated means from models across ROIs. Surface area (C), gray matter volume (D), and cortical thickness (E) expectedly follow similar patterns across regions. error bars = 95% CI

**Table 1.** Statistics for Tukey-adjusted comparisons between surface area, gray matter volume, and cortical thickness within and between ROIs.

| **Comparison** | **Estimate** | **t-ratio** | **p-value** |
| --- | --- | --- | --- |
| **Surface Area** |  |  |  |
| *LH PO – RH PO* | 272.66 | -40.32 | <0.0001 |
| *LH PT – RH PT* | -119.55 | -2.07 | 0.310 |
| *LH STG – RH STG* | 272.41 | 4.71 | 0.0001 |
| *RH PO – RH PT* | -118.52 | -2.05 | 0.320 |
| *LH PO – LH PT* | 273.69 | 4.74 | 0.0001 |
| **Gray Matter Volume** |  |  |  |
| *LH PO – RH PO* | 916 | 4.16 | 0.001 |
| *LH PT – RH PT* | -380 | -1.73 | 0.517 |
| *LH STG – RH STG* | 821 | 3.73 | 0.004 |
| *RH PO – RH PT* | -198 | -0.90 | 0.946 |
| *LH PO – LH PT* | 1098 | 4.99 | <0.0001 |
| **Cortical Thickness** |  |  |  |
| *LH PO – RH PO* | 0.03 | 1.21 | 0.834 |
| *LH PT – RH PT* | 0.02 | 0.97 | 0.925 |
| *LH STG – RH STG* | -0.004 | -0.16 | 1 |
| *RH PO – RH PT* | 0.15 | 6.33 | <0.0001 |
| *LH PO – LH PT* | 0.16 | 6.57 | <0.0001 |

*PO, pars opercularis; PT, pars triangularis; STG, superior temporal gyrus; LH, left hemisphere; RH, right hemisphere*

We next assessed whether cortical morphology was related to our behavioral outcomes (QuickSIN score, categorization gradience). Since structural MRI measures of area, volume, and thickness reflect different anatomical properties but are inherently colinear, we built separate linear regressions for each MRI measure (z-scored within-measure) per ROI. This approach resulted in 54 models (3 behavioral outcomes x 3 MRI measures x 6 ROIs). For each ROI, we applied a Holm family-wise correction for the family of regressions predicting each behavioral measure with the different anatomical metrics (e.g., QuickSIN ∼ STG_SA_; QuickSIN ∼ STG_GMV_; QuickSIN ∼ STG_CT_). This approach was taken because different behavioral measures and different ROIs addressed different questions and thus was intended to balance Type I and Type II errors.

No left hemisphere structure predicted behavioral outcomes including categorization (all *p_Holm_* > 0.248) and QuickSIN scores (all *p_Holm_* > 0.719). Surface area in RH PO (IFG) predicted categorization gradience [*F*(1, 27) = 7.27, *p* = 0.012, *p_Holm_*= 0.036, *η^2^_p_* = 0.21] (**Fig. 3A**) with more gradient listeners having greater right PO surface area. Right hemisphere STG thickness also marginally predicted categorization consistency [*F*(1, 27) = 5.43, *p* = 0.0275, *p_Holm_* = 0.082, *η^2^_p_* = 0.17] (**Fig. 3B**) with more consistent listeners having thicker right STG. Categorization consistency was not predicted by MRI measures in any other right hemisphere ROI (all other *p_Holm_* > 0.368).

**Figure 3.**
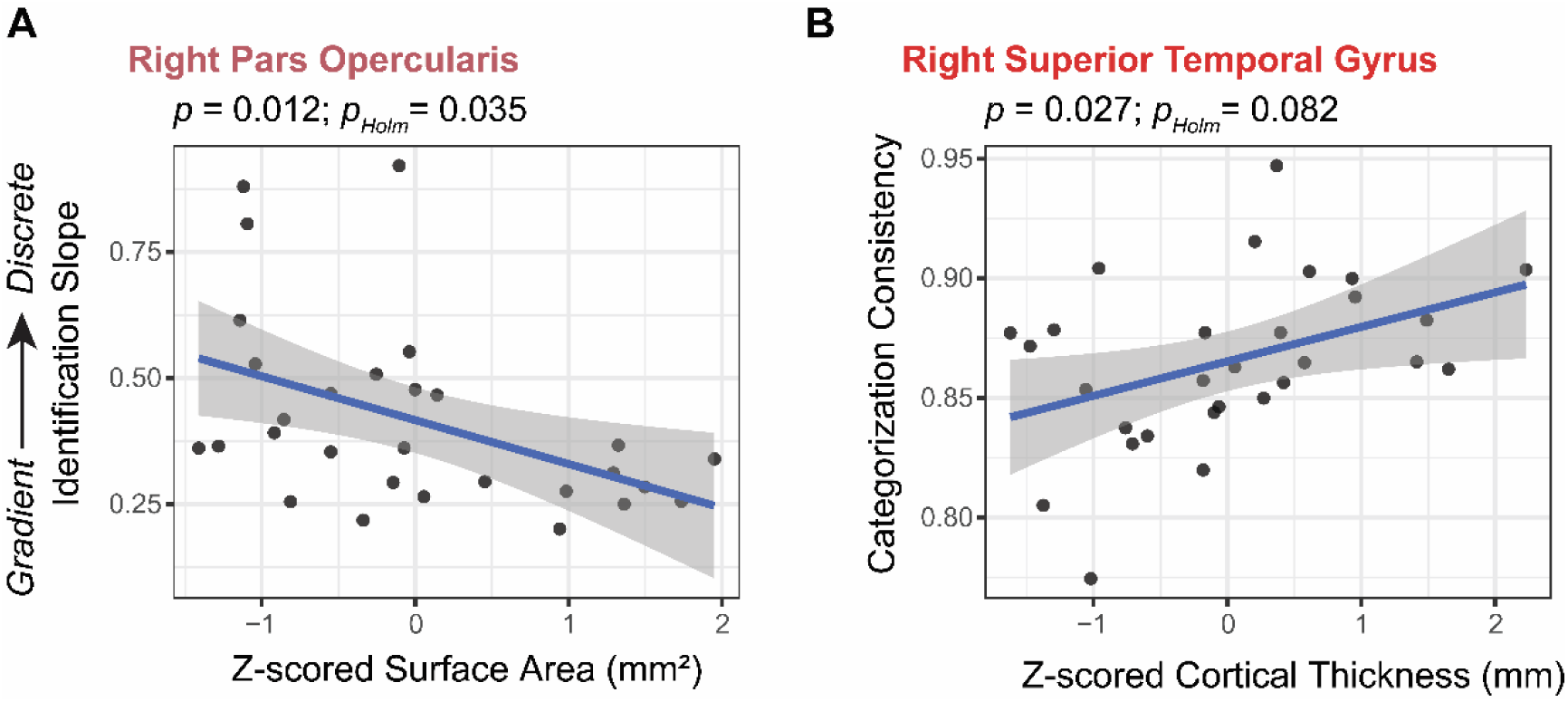
Structure-behavior relationships. (A-B) Estimated effects from models described in text with corrected and uncorrected p values shown. (A) More gradient listeners have larger surface area in RH IFG (pars opercularis). (B) More consistent listeners have greater cortical thickness in RH STG (auditory cortical region), though this effect does not remain significant when family-wise error correction is applied. Shading = 95% CI.

### 3.3 Tractography data

We next used a linear mixed effects model with a fixed effect for tract bundle (4 levels: AF_right, AF_left, BS_right, BS_left) and random intercepts for subjects [QA ∼ tract + (1|sub)] to evaluate if QA varied across tracts. There was a main effect of tract on QA [*F*(3, 87) = 154.1, *p* < 0.0001, *η^2^_p_* = 0.84] (Tukey-adjusted contrasts: all *p* < 0.007) (**Fig. 4**).

**Figure 4.**
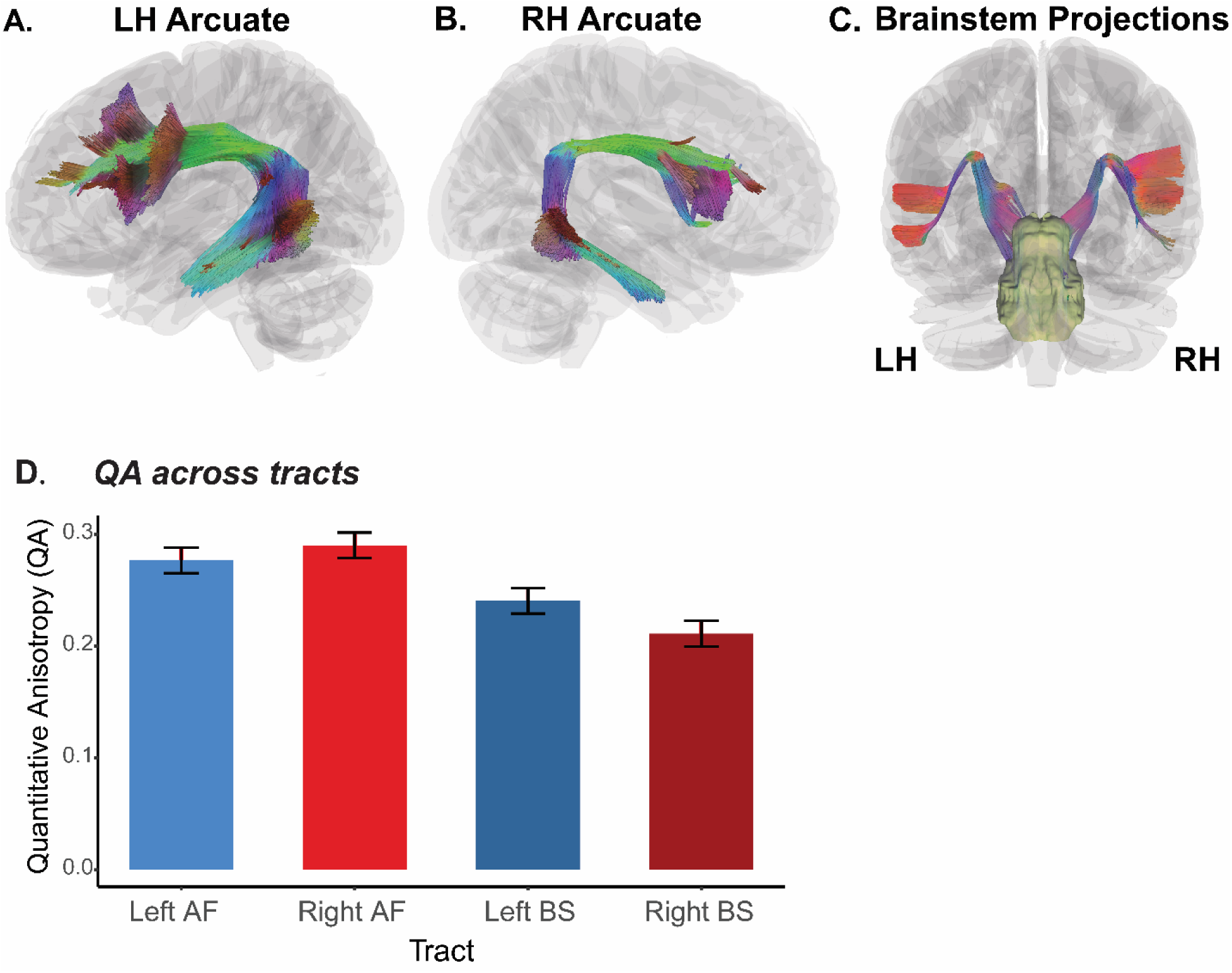
Structural connectivity results. (A-C) Grand average tractography for bilateral Arcuate Fasciculus and brainstem-cortical projections. (D) Mean QA across tracts. Right AF has the greatest fiber bundle density whereas right brainstem pathways have the lowest density. Error = s.e.m.

We next examined whether QA across tracts related to SIN performance. We used a linear fixed effects model predicting QuickSIN with fixed effects of QA from each tract across brainstem and cortical levels [QuickSIN ∼ QA_AF_Left_ + QA_AF_Right_ + QA_BS_Left_ + QA_BS_Right_].

However, variance inflation factors (VIFs: 8.0-13.9) indicated high multicollinearity among the predictors. To address this collinearity, we collapsed across hemispheres to investigate whether QuickSIN was predicted by QA at the brainstem and/or cortical level [QuickSIN ∼ QA_BS_mean_ + QA_AF_mean_]. This model reduced VIFs to >5.7, indicating more acceptable levels of multicollinearity (Kim, 2019). The model revealed that QA of the brainstem-cortical projections (but not AF) predicted QuickSIN [*F*(1, 27) = 4.45, *p* = 0.044, *η^2^_p_* = 0.14], indicating greater density in the brainstem fiber tracts predicted better (i.e., lower) QuickSIN scores (**Fig. 5B**).

**Figure 5.**
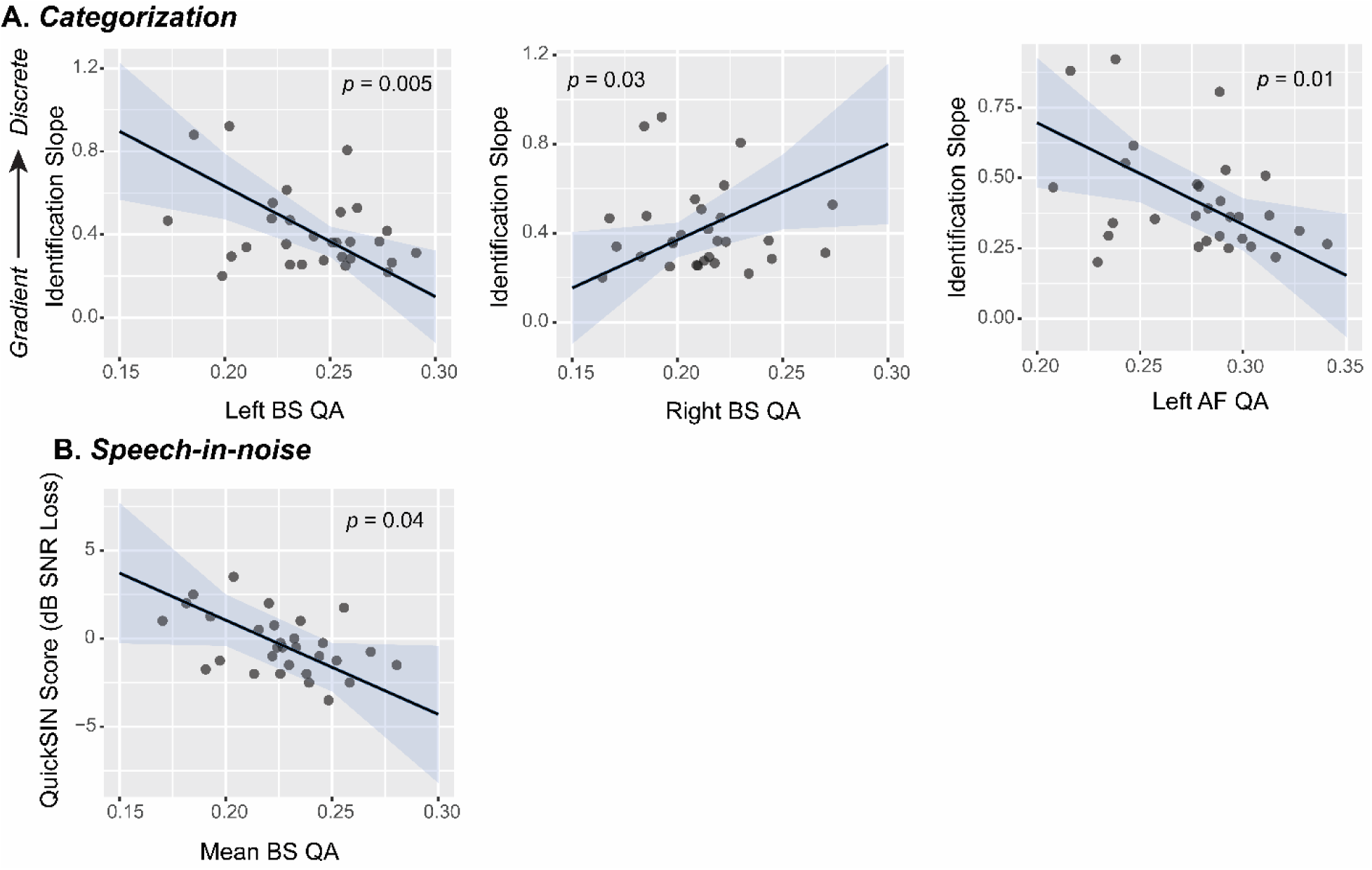
Structure-behavior relations between DWI tractography and categorization/SIN perception. (A) Categorization slope effects. White matter density (QA) shows opposite relationships to perceptual gradience across hemispheres. More gradient listeners have denser white matter in LH brainstem-cortical projections and AF, but less dense RH brainstem-cortical projections. (B) Listeners with better SIN scores have denser white matter in brainstem-cortical projections averaged across hemispheres. Shading = 95% CI.

We next investigated whether QA predicted categorization *gradience* (identification slopes). We again started with a full model predicting behavioral slope from QA at cortical and subcortical levels, as above. VIFs in this model also indicated high multicollinearity (VIF=8 to 14). To improve multicollinearity issues as well as assess possible hemispheric asymmetries, we analyzed left and right pathways separately [i.e., slope ∼ QA_BS_Left_ + QA_BS_Right_; slope ∼ QA_AF_Left_ + QA_AF_Right_]. For each of these models, VIFs remained under 2.65, indicating acceptable multicollinearity. For the brainstem model, we found significant main effects of both RH [*F*(1, 27) = 5.18, *p* = 0.031, *η^2^_p_* = 0.002] and LH [*F*(1, 27) = 9.38, *p* = 0.005, *η^2^_p_* = 0.26] QA on slope. The direction of these effects was inverse across hemispheres with a positive estimate for RH and a negative estimate for LH. That is, more gradient listeners had denser brainstem-cortical projections in the LH but sparser projections in RH (**Fig. 5A**). For the AF model, there was a significant main effect of LH QA [*F*(1, 27) = 6.93, *p* = 0.014, *η^2^_p_* = 0.20], whereby greater QA in the left arcuate predicted more gradient categorization (**Fig. 5A**).

We similarly assessed a linear regression model with QA from each tract predicting categorization *consistency*. Diagnostic VIFs ranged from 8.1 to 14.0. As with slopes, we built separate models for each anatomical level (i.e., consistency ∼ QA_BS_Left_ + QA_BS_Right_; consistency ∼ QA_AF_Left_ + QA_AF_Right_). Multicollinearity was acceptable from these models (VIF < 2.65). However, QA did not predict categorization consistency in either model (all *p* > 0.62).

## 4. Discussion

We assessed whether individual differences in speech categorization and SIN perception related to structural morphology and anatomical connectivity between major nodes of the brain’s auditory-linguistic network including the arcuate fasciculus and central auditory pathways. We found that more gradient listeners had greater surface area in right IFG, while more consistent listeners had thicker right STG, suggesting perceptual gradience and consistency may be supported by linguistic- and auditory-centric brain regions, respectively. Gradience also correlated with white matter tract density differentially across hemispheres. At a cortical level, more gradient listeners had denser white matter tracts in LH arcuate fasciculus. At a subcortical level, gradient listeners also showed denser projections of the auditory brainstem pathways in LH and sparser projection in RH compared to more discrete listeners. Denser auditory projections were further associated with better QuickSIN scores.

### 4.1 Structural Findings

#### 4.1.1 Categorization gradience and consistency relate to volumetrics of auditory-linguistic brain regions

By examining volumetric properties of listeners’ MRIs, we found that distinct auditory and linguistic brain regions related to different perceptual constructs of phonetic speech categorization. The relationship between brain structure and gradience was constrained to frontal ROIs (right PO), while that with consistency was constrained to temporal ROIs (right STG). Although entirely structural in nature, these findings are consistent with prior functional neuroimaging studies demonstrating frontal language regions, including PO, respond categorically (i.e., non-gradiently) to speech sound continua (Alho et al., 2016; Bidelman & Walker, 2019; Husain et al., 2006a; Lee et al., 2012; Luthra et al., 2019; Myers, 2007; Myers et al., 2009), as well as our recent EEG data showing perceptual consistency is functionally localized to activity from auditory cortex (Rizzi et al., 2026). That our volumetric findings were restricted to the right hemisphere is likely due to the use of vowels. Vowels are distinguished by steady-state spectral information and rely on longer integration time windows that better engage right hemispheric auditory processing (Boemio et al., 2005; Poeppel, 2003; Sininger & Bhatara, 2012; Sininger & Cone-Wesson, 2004; Zatorre & Belin, 2001). A similar rightward lateralization was observed relating fricative categorization to structural brain morphology (Fuhrmeister & Myers, 2021).

Our findings support prior work demonstrating individual differences in perceptual gradience vs. consistency might be segregated at the neuroanatomical level–supported by structural properties of frontal vs. auditory regions, respectively (Fuhrmeister & Myers, 2021). However, we hypothesized that more discrete listeners would have greater surface area in frontal regions based on findings from Fuhrmeister and Myers (2021) who demonstrated a relationship in right middle frontal gyrus (MFG). Husain et al. (2006a) also showed greater activation of MFG when categorizing nonspeech relative to speech sounds. While activation in MFG are common during tasks with categorical decisions (Mahmud et al., 2021), many studies, including our data here, emphasize the importance of adjacent IFG in speech categorization (e.g., Alho et al., 2016; Blumstein et al., 2005; Lee et al., 2012; Meyers et al., 2008; Myers et al., 2009). In right IFG (pars opercularis), we found an opposite relationship where increased surface area predicted more gradient responding. Similarly, Golestani et al. (2011) demonstrated larger left pars opercularis volume correlated with the number of years an individual had with phonetic transcription experience. Though they did not assess auditory categorization explicitly, expert phoneticians presumably have greater sensitivity to fine-grained acoustic detail (i.e., more gradience) which may allow more flexible categorization when transcribing acoustic input into an orthographic form.

We also found categorization consistency assessed under a VAS task was positively related to cortical thickness of right STG supporting our initial hypothesis that more consistent listeners have larger volumetrics in auditory cortex. Though, we note this relation did not survive family-wise error correction so the effect should be interpreted cautiously. Similarly, Fuhrmeister and Myers (2021) found less consistent listeners had increased gyrification of bilateral transverse temporal gyri (i.e., HG). When interpreted collectively with our findings, it appears consistency in categorizing speech sounds relates to structural properties of auditory cortical regions.

Though gyrification and cortical thickness are distinct measures, they are often negatively related in MRI morphometry studies in healthy middle aged adults (Gautam et al., 2015). Similarly, children with dyslexia have reduced cortical thickness and increased local gyrification in occipitotemporal regions, suggesting an inverse relationship between these metrics related to reading ability (Williams et al., 2017). Thus, the seemingly opposite relationships with consistency observed here with cortical thickness and in Fuhrmeister and Myers (2021) with local gyrification are easy to reconcile. The overall structure of auditory cortical regions, both decreased gyrification and increased cortical thickness, predict a more consistent categorizer. Gyrification of transverse HG is developed in utero and changes little with environmental factors (Golestani et al., 2011). However, cortical thickness can change with social-environmental factors such as socioeconomic status (Piccolo et al., 2016) and perceptual experience. For instance, STG thickness increases after foreign language learning (Mårtensson et al., 2012), simultaneous language interpretation training in multilinguals (Hervais-Adelman et al., 2017), and balance training (Rogge et al., 2018). Similarly, children who participated in musical training had slowed rates of cortical thinning in STG relative to controls (Habibi et al., 2020). Thus, it is possible that some listeners are more predisposed to be perceptually consistent which may relate to an anatomical morphotype with less local gyrification of transverse HG. In contrast, those who become more perceptually consistent through auditory experience have increased cortical thickness in STG. In other words, specific structural differences observed between these studies could arise from distinct origins (i.e., structural predispositions vs. experience-dependent changes in auditory perceptual skills). Our sample did not have a spread in language experience. While there was a spread in musical experience, music training was not a significant factor in any of our models when added as a covariate.

#### 4.1.2 Categorization gradience and consistency may be independent processes

The finding that gradience relates to structure of IFG while consistency relates to structure of STG suggests that these two perceptual processes of speech categorization might be subserved by distinct neural regions. Whether categorization gradience and consistency are independent attributes of behavior has been equivocal across studies (Honda et al., 2024; Kapnoula et al., 2017; Kim et al., 2025a; Myers et al., 2024; Rizzi & Bidelman, 2024, 2025); some reports show gradience is correlated with consistency in phoneme labeling while others do not. Though structural neuroimaging alone cannot discern whether perceptual processes rely on functionally distinct neural mechanisms, the anatomy of auditory cortical and brainstem regions seems to at least partially explain functional elements of auditory encoding (Bidelman et al., 2026b) and their correspondence with different anatomical regions supports the notion that they reflect independent constructs in categorization. Functionally, consistency may be more localized to auditory cortical regions (or even subcortex as suggested by Rizzi et al., 2026), whereas gradience may arise from more frontal brain areas. Prior work has shown activity in auditory cortical regions relates to the accuracy of sound identification (i.e., the quality of speech representation), while inferior frontal activity relates to the speed of identification, suggesting a functional dissociation between sensory and decision-based processes in the brain’s fronto-temporal pathways (Binder et al., 2004). If consistency arises earlier in the auditory system than gradience, gradience may be a higher-level process related to stimulus decision. That is, consistency might arise from more automatic, sensory processing. This could explain why perceptual consistency seems to be more trait-like (Kim et al., 2025b), while perceptual gradience is less stable across different phonetic contrasts (Kapnoula et al., 2021; Kim et al., 2025b; Myers et al., 2024).

### 4.2 Structural connectivity findings

#### 4.2.1 Denser WM in auditory pathways predicts better SIN understanding

DWI tractography showed that listeners with better SIN understanding had denser white matter in bilateral auditory brainstem-cortical projections that comprise the canonical central auditory system pathways. Fiber tracking does not assess functional connectivity between brain regions nor the directionality of electrophysiological signaling (only form, size, and shape of the anatomy). Thus, whether the relation between the auditory brainstem projections and SIN processing observed here is due to afferent or efferent neurophysiological function cannot be determined from our purely structural DWI data. Denser auditory pathways could support stronger and more efficient “bottom-up” encoding of speech and/or stronger efferent feedback from the descending corticofugal pathways that provide greater “top-down” control over auditory processing at lower levels. Theoretically, both the ascending and descending systems are critical for SIN processing. Robust SIN understanding requires strong bottom-up encoding to maximize the quality of sensory representation (e.g., Anderson et al., 2012; Anderson et al., 2011; Coffey et al., 2017; Parbery-Clark et al., 2009a). Indeed, reduced afferent functional connectivity between auditory midbrain and cortex correlates with poorer SIN comprehension as measured by the QuickSIN (Bidelman et al., 2019). Several electrophysiological studies in animals and humans also suggest that the corticofugal efferent system, including the cortical-brainstem connections measured here, assist in noise-degraded speech listening (Asilador & Llano, 2021; de Boer & Thornton, 2008; Price & Bidelman, 2021). Thus, the structure-behavior relationship we observe here could support either enhanced afferent or efferent functional connectivity, both of which might aid SIN perception. Further studies should systematically investigate relationships between directional brainstem-auditory tractography, functional connectivity, and SIN processing.

Contrary to our hypothesis, we did not observe a relationship between SIN performance and AF density. Although limited, prior work has suggested denser or higher integrity AF relates to better SIN perception, motivating our analysis of this pathway with DWI tractography (Li et al., 2021; Perron et al., 2021; Tremblay et al., 2019). However, these previous anatomical studies only assessed CV discrimination in noise (Li et al., 2021; Perron et al., 2021; Tremblay et al., 2019) rather than sentence-level SIN processing as used here. Given the documented role of the AF in phonological processing (Lebel & Beaulieu, 2009; Perdue et al., 2025; Tremblay et al., 2019; Vandermosten et al., 2012; Yeatman et al., 2011), it is possible sentence-level SIN perception, assessed by the QuickSIN, recruits different neural regions than simpler phonetic discrimination in noise tasks used in prior work (Li et al., 2021; Perron et al., 2021; Tremblay et al., 2019). Most studies also examined participants with extensive musical training, and musicians are known to have enhanced SIN perception (for review, see Hennessy et al., 2022; Maillard et al., 2023) and stronger AF (Halwani et al., 2011; Li et al., 2021; Moore et al., 2017; Oechslin et al., 2009; Perron et al., 2021). For example, Li et al. (2021) observed a relationship between SIN and AF microstructure but only in highly trained musicians while Perron et al. (2021) did not observe this effect in amateur singers (vs. non-singers). Similarly, while SIN advantages have been reported for musicians (e.g., Bidelman & Krishnan, 2010; Bidelman & Yoo, 2020; Hennessy et al., 2022; Maillard et al., 2023; Parbery-Clark et al., 2009b; Parbery-Clark et al., 2011; Slater et al., 2015; Yoo & Bidelman, 2019), this effect is not always observed (e.g., Boebinger et al., 2015; Madsen et al., 2019; Ruggles et al., 2014; Yeend et al., 2017). We did not explicitly recruit musicians in our study. Thus, it is possible we were unable to observe a relationship between SIN and AF given the relative homogeneity in these measures among out musically naïve listeners.

#### 4.2.2 Denser WM in LH auditory and language pathways predicts more gradient listening

With regard to anatomical correlates of categorization, we found significant asymmetries in the cortical auditory-linguistic pathways. More gradient listeners had denser AF in left hemisphere. This finding largely agrees with our hypotheses. We predicted denser WM in left hemisphere auditory and language tracts would correspond to more gradient listening. Phonetic categorization is largely a LH process with more leftward lateralized auditory and language regions specialized for phonetic category processing (Blumstein et al., 2005; Desai et al., 2008; Husain et al., 2006a; Joanisse et al., 2007; Liebenthal et al., 2005; Wolmetz et al., 2011). One study found that sensitivity on CV discrimination was related to diffusivity in bilateral AF, suggesting more developed AF in both hemispheres predicts higher sensitivity to phonetic details (Tremblay et al., 2019). Similar findings were reported by Perron et al. (2021) for left AF. Though their tasks were not canonical categorization tasks, it is possible more gradient listeners in these studies were more sensitive to the acoustic details differentiating CV contrasts (e.g., Perron et al., 2021; Tremblay et al., 2019). This could explain why gradient listening was associated with denser WM in LH auditory-language pathways. This finding also mirrors results from our recent functional (EEG) neuroimaging studies which showed that more gradient listeners had stronger left-lateralized speech activations in the temporal lobe (Rizzi & Bidelman, 2024).

The leftward asymmetry of this effect suggests that auditory perceptual gradience might depend on the dorsal speech processing stream (Hickok & Poeppel, 2004). The dorsal stream is a functional pathway that runs from left STG through the temporoparietal junction with termination in IFG. A major anatomical component of the dorsal network is the AF which is traditionally involved in sensorimotor integration and mapping sound to articulatory representations (Hickok & Poeppel, 2004, 2007). Functional neuroimaging work has suggested dorsal stream involvement in speech categorization (Alho et al., 2016; Chevillet et al., 2013; Lee et al., 2012). We extend these prior results by establishing a structural basis for such effects. Moreover, our findings lead us to infer that the dorsal stream not only contributes to phonetic categorization very broadly, but more importantly, to how nuanced (gradient) a listener maps speech sounds to their identity.

Notably, we also found structural asymmetries within the auditory brainstem-cortical pathways. Though some studies have tracked changes in the central auditory brainstem pathways in relation to hearing acuity and tinnitus (Koops et al., 2021; Svobodová et al., 2024), none to our knowledge have explored whether their density maps to perceptual processing. Here, we show that more gradient listeners have denser left but sparser right WM in their auditory tracts, supporting our hypothesis that more continuous modes of perception would relate to auditory system neuroanatomy. Presumably, denser WM in auditory system may allow for more robust auditory encoding and precise capture of fine-grained acoustic details for later perceptual judgements. Interestingly, the lateralization of the brainstem data is also internally consistent with the cortical results: more gradient listeners have denser WM along the left auditory brainstem pathways that persists into left lateralized language circuitry at the cortical level.

## 5. Conclusions

We measured volumetrics (MRI) and structural connectivity (DWI) within and between major auditory and language brain and evaluated their relationship to auditory-perceptual categorization and SIN abilities. Structurally, we found gradience and consistency in speech-sound labeling related to volumetric measures in distinct brain regions; increased gradience predicted by greater surface area in right IFG and increased consistency predicted by thicker right STG. These anatomical findings imply that a consistent perceptual readout of the speech signal may be more automatic and supported by auditory cortical regions, while gradience may arise from higher-order frontal regions, which could be more influenced by attentional processing.

Structural tractography demonstrated denser WM in the auditory system was related to both improved SIN perception and categorization gradience, emphasizing the role of central auditory pathway integrity for multiple auditory perceptual skills. Specifically, increased gradience was predicted by denser left-lateralized WM in both the auditory and language tracts, suggesting these LH tracts may play a role in retaining subphonemic detail up to the level of IFG for phonetic categorization.

This work expands on our prior *functional* EEG experiments (Rizzi & Bidelman, 2024; Rizzi et al., 2026) by revealing *structural*-behavioral relationships that also underly individual differences in listening strategies as they relate to speech categorization. The MRI/DWI results herein highlight the importance of the central auditory system pathways and dorsal speech processing stream in accounting for gradience and consistency in auditory category mapping. That brain-behavior relationships were largely left lateralized across subcortical and cortical fiber tracts implies an integrated network of auditory and language pathways that underly phonetic categorization gradience.

## Acknowledgements

The authors thank Tessa Bent, Jennifer Lentz, and Samantha Gustafson for their comments on earlier versions of this manuscript.

## Author contributions

R. R. and G.M.B. designed the experiment, R.R., J.R.S., and Z.E. collected the data, R.R. and G.M.B. analyzed the data, and all authors wrote the paper.

## Funding

This work was supported by the National Institutes of Health/National Institute on Deafness and Other Communication Disorders R01DC016267 (awarded to G. M. B.) and F31DC023124 (awarded to R. R.). This project was also supported by the Indiana Clinical and Translational Sciences Institute (CTSI), funded in part by grant #UL1TR002529 from the National Institutes of Health, National Center for Advancing Translational Sciences, Clinical and Translational Sciences Award.

## Data Availability

The datasets generated during the current study are available from the corresponding author on reasonable request.

## Conflict of interest

The authors have no relevant financial or non-financial interests to disclose.

## Ethical approval

This study was performed in line with the principles of the Declaration of Helsinki. Approval was granted by the Institutional Review Board of Indiana University (No. 15650, approved 12/9/25).

## Notes

### Competing Interest Statement

The authors have declared no competing interest.

